# NanoCellAnnotator: Formalizing Expert Cell Type Annotation with Large Language Models

**DOI:** 10.64898/2026.06.21.728965

**Authors:** Md Ishtyaq Mahmud, Veena Kochat, Humaira Anzum, Suresh Satpati, Jagan Mohan Reddy Dwarampudi, Kunal Rai, Tania Banerjee

## Abstract

**Motivation:** Cell-type annotation in spatial transcriptomics is challenging due to sparse gene panels, spatial heterogeneity, and limited availability of tissue-matched reference atlases. Recent approaches have explored large language models (LLMs) for integrating biological knowledge during annotation, but unconstrained inference can produce biologically unsupported predictions and hallucinated cell types. In addition, many LLM-based pipelines rely on large cloud-hosted models that limit reproducibility and deployment in privacy-sensitive environments.

**Results:** We introduce *NanoCellAnnotator*, a biologically constrained and confidence-aware framework for automated cell-type annotation in spatial transcriptomics. The framework decouples spatial structure discovery, deterministic biological evidence construction, and language-model-based semantic inference. Spatial clusters are identified using hybrid spatially regularized non-negative matrix factorization (hSNMF), after which cluster-level marker genes are abstracted into ontology-derived functional programs using Gene Ontology enrichment and GO-slim projection. A lightweight locally executable language model performs constrained label selection within a curated admissible label space derived from PanglaoDB and CellMarker. Annotation confidence is estimated independently using marker support strength and lineage separation, enabling ambiguous or heterogeneous clusters to be explicitly flagged. We evaluate NanoCellAnnotator on Xenium spatial transcriptomics data from intrahepatic cholangiocarcinoma and an independent breast cancer spatial transcriptomics dataset. The framework recovers canonical cell populations with high confidence while identifying heterogeneous or transitional spatial domains as ambiguous. Agreement with manual annotations was evaluated using accuracy and adjusted Rand index.

**Availability:** Code available at https://github.com/ishtyaqmahmud/NanoCellAnnotator.

## 1 Introduction

Spatial transcriptomics technologies enable the measurement of gene expression while preserving spatial context within intact tissue, providing new opportunities to study tissue organization, cellular interactions, and disease-associated microenvironments. Recent platforms such as 10x Genomics Xenium provide single-cell resolution measurements over targeted gene panels, enabling spatial profiling across large tissue sections [1].

Accurate cell-type annotation remains a central challenge in spatial transcriptomics. Many existing annotation methods were originally designed for single-cell RNA sequencing and rely on supervised classifiers or reference-based label transfer. These approaches assume dense gene coverage and the availability of matched reference atlases, assumptions that often do not hold in spatial assays where gene panels are sparse and tissue-specific reference data may be unavailable.

Large language models (LLMs) have recently been explored as tools for integrating heterogeneous biological knowledge during cell-type annotation. By mapping molecular evidence to semantic cell-type concepts, LLMs can provide flexible reasoning over marker genes and biological pathways. However, unconstrained language-model inference can generate biologically unsupported predictions or hallucinated cell identities [2], [3]. Furthermore, many LLM-based annotation pipelines depend on large cloud-hosted models, limiting reproducibility and practical deployment in clinical or privacy-sensitive settings [4] [5].

Another important challenge arises from the spatial nature of the data itself. Spatial clusters frequently represent heterogeneous cellular neighborhoods or transitional states rather than pure cell populations [6], [7]. Forcing such clusters into a single canonical cell-type label can obscure biologically meaningful uncertainty.

To address these challenges, we propose *NanoCellAnnotator*, a biologically constrained and confidence-aware framework for spatial transcriptomics annotation. The key idea is to separate spatial structure discovery, deterministic biological evidence construction, and semantic inference into distinct stages. Rather than allowing an LLM to freely generate labels, the model performs constrained label selection within a curated admissible label space derived from established marker databases. Annotation confidence is estimated independently using marker-gene support and lineage separation, allowing ambiguous clusters to be explicitly flagged rather than forcibly labeled.

We demonstrate the framework on spatial transcriptomics datasets from intrahepatic cholangiocarcinoma and breast cancer. Results show that biologically constrained language-model inference enables reliable and interpretable annotation while preserving uncertainty in heterogeneous spatial domains.

## 2 Methods

### 2.1 Overview of the NanoCellAnnotator Framework

NanoCellAnnotator consists of three stages: spatial clustering, biological evidence construction, and constrained semantic inference using a lightweight language model.

While recent large language model (LLM)–based approaches have shown promise for cell-type annotation in single-cell RNA-sequencing data, their direct application to spatial transcriptomics is complicated by increased data sparsity, spatial heterogeneity, and the risk of biologically unsupported predictions. NanoCellAnnotator addresses these challenges by *decoupling spatial domain identification and biological evidence construction from language-model inference*. Spatially coherent cellular domains are identified upstream using spatially regularized clustering, and all downstream reasoning is performed at the level of cluster-specific abstractions rather than individual cells.

The resulting structured representations, namely, cluster-level marker genes, ontology-derived functional programs, and quantitative expression statistics, are assembled into a standardized interface that encodes an explicit hierarchy of biological evidence. A lightweight base language model is then employed solely as a constrained semantic integrator to select a single plausible cell-type label per cluster, without access to raw expression matrices, spatial coordinates, or external atlases. Finally, annotation confidence is assessed independently of the language model using marker-gene support strength and lineage separation, allowing clusters with insufficient or conflicting evidence to be explicitly flagged as ambiguous rather than forcibly labeled.

In this study, NanoCellAnnotator is evaluated on 10x Genomics Xenium spatial transcriptomics data from human bile-duct tissue. Figure 1 provides a schematic overview of the framework, highlighting the strict separation between upstream spatial analysis, deterministic biological abstraction, constrained language-model inference, and confidence-aware annotation.

**Figure 1.**
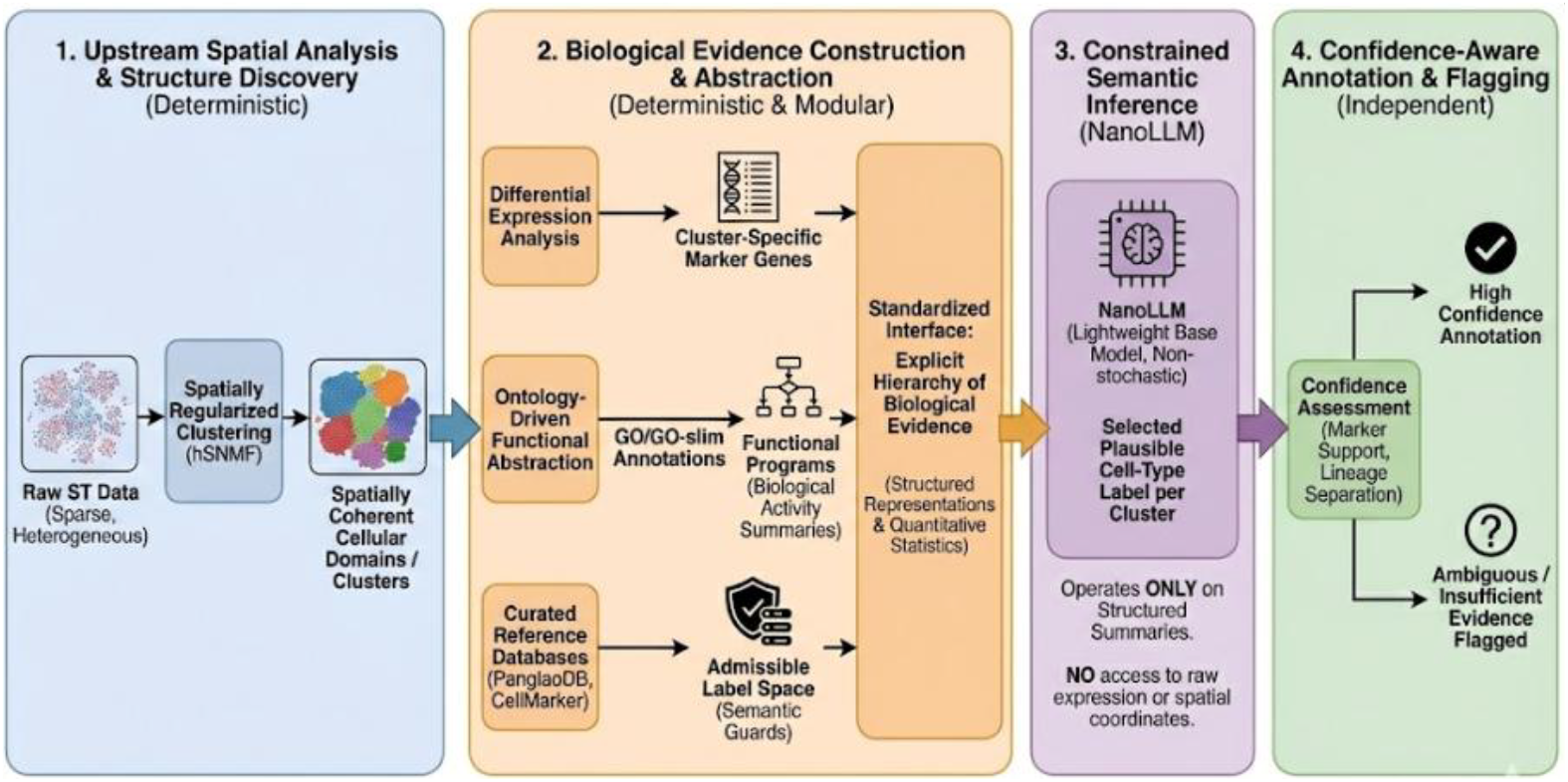
Schematic of the NanoCellAnnotator framework. The pipeline decouples spatial clustering from semantic inference to ensure reproducibility. (1) Spatial Structure Discovery: Raw data is processed via hybrid spatially regularized NMF (hSNMF) to identify coherent tissue domains. (2) Biological Evidence Construction: Marker genes are mapped to functional GO terms and constrained by reference databases (PanglaoDB, CellMarker). (3) Constrained Semantic Inference: A lightweight LLM (NanoLLM) assigns labels using only structured summaries (without access to raw expression) to ensure deterministic outputs. (4) Confidence Assessment: Final annotations are validated against marker support to explicitly flag ambiguous clusters.

**Figure 2.**
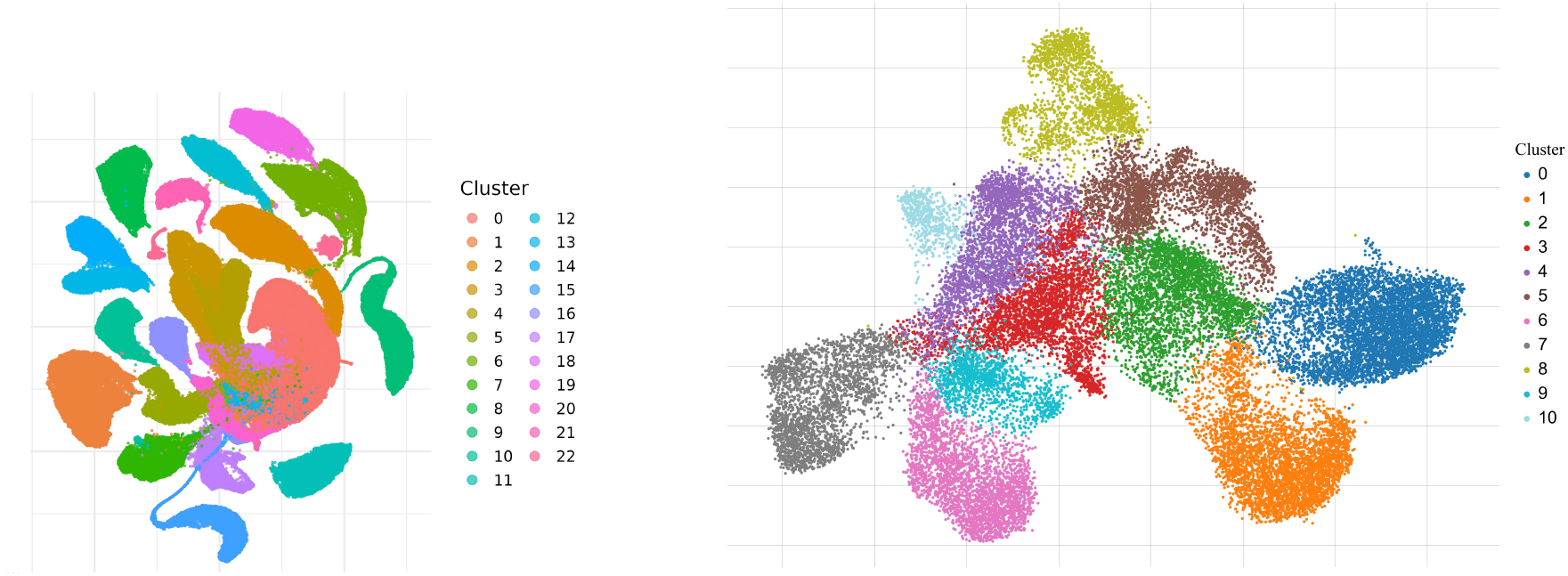
Clustering results across spatial transcriptomics datasets. **(left)** Cholangiocarcinoma (CCA) spatial clusters obtained using NMF-based dimensionality reduction with spatial regularization (hSNMF) followed by Leiden clustering. **(right)** Breast cancer (BRCA) dataset clustered using PCA for dimensionality reduction followed by Leiden clustering.

**Figure 3.**
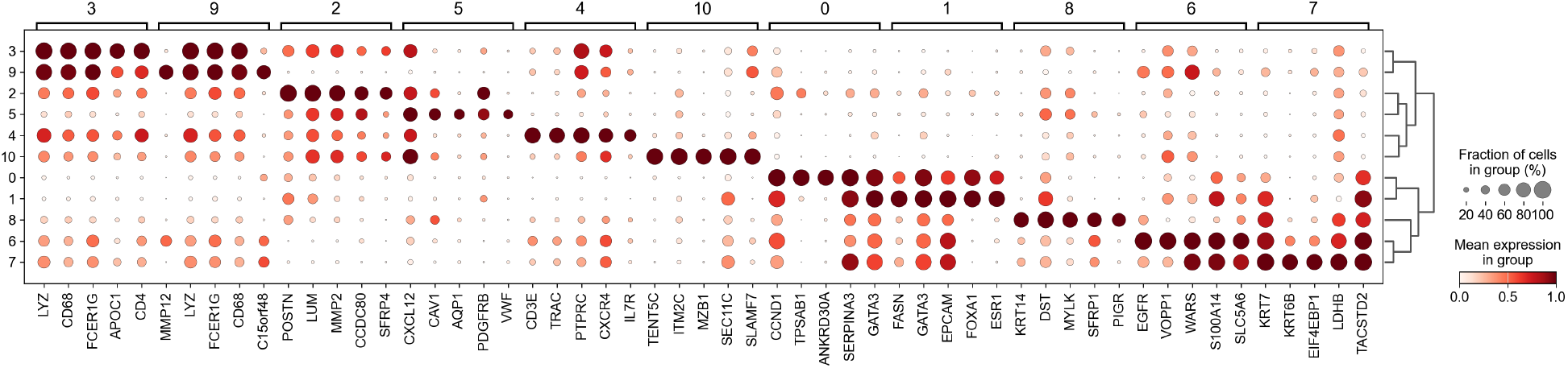
Spatially regularized biological validation. **a)** Marker gene dot plot for CCA clusters, showing distinct transcriptional signatures. **(b)** Marker gene dot plot for BRCA dataset, confirming robust biological separation.

### 2.2 Spatial Clustering and Cluster Marker Gene

To ensure that downstream cell-type annotation operates on biologically meaningful and spatially coherent units, NanoCellAnnotator performs all semantic inference at the level of spatial clusters rather than individual cells. Spatial clustering and marker gene identification are therefore treated as an upstream, model-agnostic preprocessing stage whose sole role is to define coherent cellular domains and their associated molecular signatures. All subsequent biological abstraction and language-model inference is constrained to these cluster-level representations.

#### 2.2.1 Spatially Regularized Clustering with hSNMF

To identify spatially coherent cellular domains within high-dimensional Xenium spatial transcriptomics data, we employed *Hybrid Spatially Regularized Non-negative Matrix Factorization* (hSNMF) [8] as the upstream clustering strategy. Unlike conventional dimensionality reduction methods that treat cells as independent observations, hSNMF explicitly incorporates spatial topology to promote the discovery of geometrically contiguous and transcriptionally coherent domains. Although we use hSNMF in this study, NanoCellAnnotator is agnostic to the specific clustering method and can operate on any spatially coherent clustering of cells.

Given the log-normalized gene expression matrix *X* ∈ *R*^n×m^, where *n* denotes the number of cells and *m* the number of measured genes, hSNMF computes a low-dimensional nonnegative factorization *X* ≈ *WH*. The latent representations encoded in *W* capture shared transcriptional structure while mitigating noise inherent in high-dimensional expression space.

Spatial coherence is enforced through iterative diffusion of latent representations over a spatial adjacency graph, smoothing factor values across neighboring cells while preserving underlying transcriptional variation. Importantly, spatial diffusion is applied exclusively to the cell-wise latent representation matrix *W*, while the gene-loading matrix *H* remains fixed; no spatial smoothing is performed over gene loadings or gene–gene relationships. To integrate both local cell–cell interactions and broader tissue context, we construct a hybrid spatial graph combining short-range contact-based edges (*r*_*c*_ = 20 *µ*m) with longer-range radius-based connections (*r*_*r*_ = 80 *µ*m). These spatial relationships are encoded in a weighted adjacency matrix.

Final cluster assignments are obtained by applying Leiden clustering to a mixed adjacency matrix that linearly interpolates between transcriptomic similarity and spatial proximity. Although hSNMF produces spatially smoothed latent representations, these embeddings are used exclusively for graph construction and clustering. Only the resulting discrete cluster assignments are propagated to downstream analysis; latent factors and spatial coordinates are not exposed to subsequent stages of NanoCellAnnotator.

This design ensures that all downstream biological abstraction and semantic inference operate on spatially coherent and transcriptionally meaningful cellular domains, while preventing direct reliance on latent embeddings or spatial information beyond the clustering stage. Clustering is not optimized for downstream cell-type labels and does not use reference annotations or ontological information.

In summary, hSNMF first learns expression-based cell representations, then smooths those representations across spatial neighbors, and finally clusters cells using a balanced combination of spatial proximity and transcriptional similarity.

#### 2.2.2 Cluster-Specific Marker Gene Identification

Following spatial clustering, we performed cluster-wise differential expression analysis to identify marker genes characterizing each spatial domain. For each cluster *C*_*i*_, gene expression within the cluster was compared against all other cells, and genes were ranked based on relative overexpression, statistical significance, and expression specificity. All clustering and marker identification steps are deterministic given fixed parameters.

We selected the top *k* = 20 marker genes per cluster, a fixed choice applied uniformly across all clusters. This value reflects a balance between capturing sufficient biological signal and maintaining interpretability given the fixed 480-gene Xenium panel. The resulting marker sets provide a compact, high-signal molecular summary of each spatial domain and serve as the sole gene-level input for all subsequent stages of NanoCellAnnotator.

Importantly, no raw expression values, latent embeddings, or spatial coordinates are provided to downstream functional abstraction or language-model inference. All subsequent reasoning operates exclusively on these standardized cluster-level marker sets, ensuring interpretability, reproducibility, and consistent treatment of spatial domains.

### 2.3 Knowledge Retrieval and Ontology-Based Functional Abstraction

To bridge cluster-level marker genes with biologically grounded semantic inference, NanoCellAnnotator incorporates a structured knowledge grounding and functional abstraction stage. This stage constructs all biological evidence used downstream while explicitly avoiding direct cell-type assignment or model-driven interpretation. All operations are deterministic and performed independently of the language model.

#### 2.3.1 Curated Reference Databases for Constrained Inference

NanoCellAnnotator leverages curated reference databases exclusively to constrain the admissible space of cell-type labels during semantic inference. Specifically, PanglaoDB and CellMarker are used as complementary resources providing curated associations between genes and established cell types across tissues and lineages.

Importantly, these databases are *never used* to derive gene-level biological functions, pathway annotations, or functional programs. Their role is strictly limited to acting as semantic guards that restrict the label space available to the downstream language-model inference module. For each spatial cluster, candidate cell-type labels are limited to those for which at least one cluster marker gene has documented support in PanglaoDB or CellMarker. Cell-type labels lacking curated marker evidence are excluded prior to inference.

By decoupling semantic constraints from biological inter-pretation, this design prevents unsupported or speculative annotations while preserving flexibility to reason over heterogeneous molecular evidence.

### 2.4 Ontology-Based Functional Abstraction

Cluster marker genes are summarized using Gene Ontology (GO) enrichment analysis restricted to Biological Process terms. Significant terms are projected onto a GO-slim ontology to produce compact functional summaries describing coordinated biological activity within each cluster. These functional programs provide contextual biological evidence but are not used to directly assign cell-type labels. Instead, they serve as structured inputs to the constrained language-model inference stage.

#### 2.4.1 Structured Evidence Representation

For each spatial cluster, NanoCellAnnotator constructs a structured representation comprising: (i) cluster-specific marker genes, which serve as the primary molecular evidence; (ii) ontology-derived GO-slim functional programs with their supporting genes, which provide contextual biological abstraction; and (iii) quantitative differential expression statistics, including effect size and expression specificity.

This representation encodes an explicit hierarchy of biological evidence in which marker genes constitute the irreducible basis for inference, while functional programs summarize coordinated biological activity without imposing cell-type assumptions. The resulting structured summaries form the sole input to the constrained language-model inference stage, ensuring that downstream semantic inference integrates empirical expression evidence with ontology-grounded biological context in a transparent and reproducible manner.

### 2.5 NanoLLM-Guided Cell-Type Annotation

NanoCellAnnotator employs a lightweight language model solely as a constrained semantic inference component, operating on structured biological summaries rather than raw gene expression data, spatial coordinates, or latent embeddings. Cell-type annotation is formulated as a bounded label selection task rather than a generative prediction problem.

For each spatially coherent cluster, inference is performed using Qwen2.5-1.5B-Instruct executed locally to ensure reproducibility and transparency. The model receives a structured summary of pre-computed biological evidence consisting of: (i) cluster-specific marker genes identified via differential expression analysis, (ii) ontology-derived GO-slim functional programs summarizing coordinated biological activity, and (iii) a restricted set of admissible cell-type labels defined by curated reference databases. The NanoLLM does not receive confidence scores, marker support values, candidate label rankings, or acceptance thresholds. Consequently, it cannot condition its predictions on downstream confidence-based ambiguity handling and has no influence over whether its predicted label is ultimately accepted, downgraded, or flagged as ambiguous.

The language model is constrained to select a label exclusively from this admissible set and is not permitted to generate novel or unsupported cell-type identities.

Inference is performed using deterministic greedy decoding with fixed generation parameters. The model is instructed to output exactly one concise cell-type label per cluster, without free-form explanation or multi-label output. This configuration eliminates stochastic variability and minimizes sensitivity to prompt formulation, yielding consistent and reproducible annotations across runs.

For each cluster, NanoCellAnnotator reports the inferred cell-type label together with its supporting marker genes, associated GO-slim functional programs, and a quantitative confidence score. Confidence estimation is performed independently of the language model based on marker enrichment strength and lineage purity, allowing clusters with insufficient or conflicting evidence to be explicitly flagged as ambiguous rather than forcibly annotated.

Overall, this design ensures that language-model–assisted annotation in NanoCellAnnotator remains biologically grounded, reproducible, and interpretable, with the language model functioning strictly as a constrained semantic integrator rather than a source of novel biological claims.

### 2.6 Hallucination Control via Curated Constraints and Evaluation Metrics

Unlike post hoc filtering strategies that discard or relabel predictions after inference, hallucination control is integrated directly into the annotation pipeline through constrained inference and independent confidence assessment.

Unconstrained language-model inference is susceptible to hallucinated or biologically unsupported predictions, particularly in complex tissues where marker genes may be shared across multiple lineages. Rather than relying on post-hoc validation against external atlases, NanoCellAnnotator mitigates hallucinations through a combination of *pre-inference semantic constraints* and *quantitative confidence assessment*.

#### 2.6.1 Curated Constraint of the Label Space

To prevent unsupported cell-type predictions, our annotator restricts the set of admissible cell-type labels prior to inference using curated reference databases. Specifically, PanglaoDB and CellMarker are used to define associations between genes and established cell types. For each cluster, candidate cell-type labels are limited to those for which at least one cluster marker gene is supported by curated evidence in these databases. Labels lacking documented marker support are excluded from consideration.

This design ensures that the language model is never exposed to an unrestricted vocabulary of possible cell identities and cannot generate novel or speculative labels. Importantly, curated databases are used solely to constrain the label space and do not contribute functional annotations or pathway information.

#### 2.6.2 Confidence-Based Ambiguity Handling

Even under a constrained label space, spatial clusters may exhibit overlapping marker signatures or insufficient separation between competing lineage hypotheses. To explicitly account for this uncertainty, NanoCellAnnotator computes a cluster-level confidence score based on marker-gene evidence and lineage separation, as summarized in (Algorithm 1, Supplementary Material).

Confidence estimation is performed independently of language-model inference. Scores are computed solely from cluster-specific marker genes using differential expression effect size, expression specificity, and statistical significance, and do not depend on the label predicted by the language model. The role of confidence scoring is therefore to determine whether the molecular evidence is sufficient to support assigning any definitive cell-type label.

For each cluster, candidate label support scores are obtained by aggregating weighted contributions from marker genes mapped to curated cell-type labels. Individual marker contributions are weighted by effect size, expression specificity, and statistical significance, with effect size providing the dominant signal. Support scores are summed across genes to yield a raw score for each candidate label.

Lineage separation and internal consistency are assessed by comparing the top-ranked label against competing hypotheses. Specifically, we compute (i) a relative margin between the top two candidate support scores and (ii) the Shannon entropy of the normalized candidate score distribution. These diagnostics capture whether evidence is concentrated on a single lineage or diffusely distributed across multiple labels.

The final confidence score integrates normalized top-label support, separation margin, mapping coverage, and an entropy-based consistency penalty into a bounded scalar in [0, 1]. Clusters are assigned to confidence tiers using fixed thresholds: scores ≥ 0.70 are designated as high confidence, scores between 0.50 and 0.70 as moderate confidence, and scores below 0.50 as low confidence. Clusters failing minimum evidence criteria, including insufficient marker-to-label mapping coverage or weak separation between competing labels, are flagged as ambiguous or unresolved. All thresholds and weighting parameters were fixed *a priori* and applied uniformly across all clusters.

##### Top-*k* Evidence Agreement Metric

To audit NanoLLM label selection independently of clustering quality, we define a top-*k* evidence agreement metric based on curated marker– cell-type associations. For each spatial cluster *C*, candidate cell-type labels are generated using PanglaoDB [9] and CellMarker [10] and assigned support scores by aggregating marker-level evidence contributions as defined in Algorithm 1. Candidate labels are ranked in descending order of support. Unlike accuracy-based metrics that require ground-truth labels, top-k evidence agreement measures whether the language model’s prediction is consistent with independently aggregated molecular evidence. A top-k match indicates that the predicted label is among the most strongly supported biological hypotheses for that cluster, even when fine-grained subtype resolution is ambiguous. This metric therefore evaluates semantic alignment with evidence rather than correctness relative to an external reference.

To prevent nonspecific categories from dominating evaluation, a fixed set of generic labels (e.g., “normal cell”, “unknown”) is removed prior to ranking. Let *C*_*k*_(*C*) denote the top-*k* remaining candidate labels for cluster *C* (or all candidates if fewer than *k* remain). The NanoLLM prediction 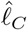 is considered a match at rank *k* if 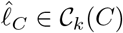.

Top-*k* agreement is reported as a cell-weighted fraction across clusters:

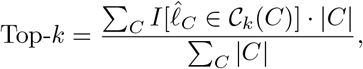

where |*C*| denotes the number of cells in cluster *C*.

## 3 Results

### 3.1 Datasets

We analyzed 10x Genomics Xenium spatial transcriptomics data from 40 tissue microarray cores obtained from 25 patients with intrahepatic cholangiocarcinoma, comprising approximately 212,000 spatially resolved cells measured over a fixed 480-gene panel. Standard quality control, normalization, and preprocessing steps were applied, including filtering low-quality cells and genes, doublet removal, and log-normalization. After preprocessing, 191,125 high-confidence single cells were retained for downstream analysis.

### 3.2 Public Dataset

We analyzed a comprehensive 10x Genomics Xenium spatial transcriptomics dataset, which was subsetted to isolate a targeted breast cancer tissue microarray (tTMA1) core initially containing 30,917 cells measured across a 541-gene panel [11]. Rigorous quality control protocols were then applied to eliminate segmentation artifacts and optical noise, including filtering out 106 low-quality cells exhibiting fewer than 10 total transcript counts. The resulting high-confidence dataset retained 30,811 cells structurally integrated with critical topological metadata, such as exact spatial coordinates, cell area, and nucleus area. This highly refined, spatially resolved feature matrix provides an optimized, low-noise foundation for downstream deep learning applications and complex neighborhood modeling.

## 4 Results

### 4.1 Spatial Clustering and Annotation Overview

Application of NanoCellAnnotator to the Xenium bile-duct dataset identified 23 spatially coherent cellular clusters representing distinct transcriptional domains. Cluster sizes ranged from 1,886 to 29,921 cells, with a median size of approximately 7,900 cells.

For each cluster, NanoCellAnnotator produced a candidate cell-type annotation together with a confidence score derived from marker-gene support and lineage separation. Based on this score, clusters were categorized into three confidence tiers: high, moderate, and low confidence.

Across the dataset, 10 clusters were assigned high confidence, 8 clusters moderate confidence, and 5 clusters low confidence. High-confidence clusters corresponded to canonical cell types such as hepatocytes, epithelial cells, cholangiocytes, and immune populations with well-supported marker signatures. Clusters assigned lower confidence typically exhibited weaker or overlapping marker evidence across candidate cell types.

A summary of cluster sizes, predicted labels, and confidence tiers is shown in Table 1.

**Table 1.** Cluster-level NanoCellAnnotator predictions for the CCA dataset.

| Cluster | Size | NanoLLM predicted label | Tier | Confidence |
| --- | --- | --- | --- | --- |
| 1 | 29921 | T Cell | High | 0.71 |
| 2 | 16679 | Mesenchymal progenitors | Moderate | 0.57 |
| 3 | 12309 | Adipocyte | Moderate | 0.59 |
| 4 | 10522 | Cholangiocyte (Reactive/EMT-like) | High | 0.89 |
| 5 | 10090 | Epithelial cells | Low | 0.35 |
| 6 | 9271 | Hepatocyte | High | 0.84 |
| 7 | 8775 | Cholangiocyte (Reactive/EMT-like) | Low | 0.40 |
| 8 | 8211 | Hepatocyte | High | 0.78 |
| 9 | 7960 | Epithelial cells | High | 0.89 |
| 10 | 7906 | Hepatocyte | High | 0.85 |
| 11 | 7862 | Epithelial cells | High | 0.71 |
| 12 | 7384 | Hepatocyte | Moderate | 0.63 |
| 13 | 6896 | Cholangiocyte (Tumor) | High | 0.89 |
| 14 | 6292 | Hepatocyte | Moderate | 0.59 |
| 15 | 5767 | Hepatocyte | Moderate | 0.52 |
| 16 | 5400 | Hepatocyte | Moderate | 0.57 |
| 17 | 5396 | Epithelial cells | Moderate | 0.65 |
| 18 | 4975 | Cholangiocyte (Reactive/EMT-like) | High | 0.90 |
| 19 | 4926 | Liver cell | Low | 0.46 |
| 20 | 4757 | Hepatocyte | Moderate | 0.60 |
| 21 | 4217 | Cholangiocyte (Reactive/EMT-like) | High | 0.90 |
| 22 | 3723 | Mesenchymal progenitors | Moderate | 0.57 |
| 23 | 1886 | Fibroblast / Stroma | Low | 0.41 |

### 4.2 BRCA Dataset Annotation and Agreement with Reference Labels

To evaluate annotation accuracy on an independent dataset, NanoCellAnnotator was applied to a breast cancer (BRCA) spatial transcriptomics dataset with manually curated reference annotations.

The pipeline identified 11 clusters ranging in size from 679 to 5,306 cells. Table 3 summarizes NanoCellAnnotator predictions together with the corresponding manual annotations.

Agreement between predicted and manual annotations was 48.2% at the cell-type level and 56.8% at the lineage level. Most disagreements occurred in tumor-associated clusters, where transcriptional programs may not correspond directly to canonical immune or stromal cell-type labels.

### 4.3 Confidence-Aware Cell-Type Annotation

Clusters assigned high confidence by NanoCellAnnotator corresponded to well-established cellular identities with strong marker-gene support (Table 1 and 2). In contrast, clusters assigned lower confidence typically exhibited weaker or overlapping marker signatures across multiple candidate cell types. These results indicate that the confidence score reflects the strength and specificity of the underlying molecular evidence.

**Table 2.** Cluster-level NanoCellAnnotator predictions and confidence scores for the BRCA dataset.

| Cluster | Size | NanoLLM predicted label | Tier | Confidence |
| --- | --- | --- | --- | --- |
| 1 | 5306 | Tumor Luminal Proliferative | High | 0.69 |
| 2 | 4001 | Tumor Luminal Proliferative | Low | 0.50 |
| 3 | 3175 | Macrophage | Moderate | 0.55 |
| 4 | 3053 | Tumor Luminal Proliferative | High | 0.78 |
| 5 | 3016 | T cells | High | 0.79 |
| 6 | 2853 | Tumor Luminal Proliferative | High | 0.90 |
| 7 | 2845 | Mesenchymal Progenitors | High | 0.73 |
| 8 | 2539 | Tumor Basal Active | High | 0.73 |
| 9 | 1966 | Fibroblast | High | 0.68 |
| 10 | 1378 | Tumor Luminal Proliferative | High | 0.86 |
| 11 | 679 | Macrophage | High | 0.80 |

### 4.4 Ablation Study

We performed ablation analyses to quantify how curated label constraints and structured biological evidence affect the behavior of the semantic inference stage. These experiments characterize the contribution of individual components of NanoCellAnnotator and complement evaluation against external reference annotations.

Ablation effects are evaluated using a *top-k evidence agreement* metric, defined as the cell-weighted fraction of clusters for which the NanoLLM prediction appears among the top-*k* candidate cell-type labels derived from independent marker-based evidence. To prevent nonspecific database categories from dominating evaluation, generic labels (e.g., “normal cell”, “unknown”) were excluded from candidate ranking prior to computing agreement.

All ablation metrics are computed on clusters with *High* or *Moderate* cluster-level confidence, as determined by independent marker-evidence scoring. Clusters flagged as *Low* confidence were excluded from NanoLLM evaluation, yielding a cell-weighted commitment rate of 86.57%. Unless stated otherwise, A1 (full evidence with admissible label constraints) serves as the canonical reference configuration for all ablation comparisons.

We evaluate six ablation configurations that selectively vary the evidence provided to the NanoLLM and the application of admissible label constraints (Table 4). The remaining two possible combinations were not evaluated because they do not correspond to meaningful inference settings within the NanoCellAnnotator pipeline.

**Table 3.** NanoCellAnnotator predictions compared with manual annotations for the BRCA dataset.

| Cluster | Size | NanoLLM predicted label | Manual Annotation |
| --- | --- | --- | --- |
| 1 | 5306 | Tumor Luminal Proliferative | Tumor Luminal Proliferative |
| 2 | 4001 | Tumor Luminal Proliferative | Tumor Luminal Proliferative |
| 3 | 3175 | Macrophage | Fibroblast |
| 4 | 3053 | Tumor Luminal Proliferative | Macrophage |
| 5 | 3016 | T cells | T Cell |
| 6 | 2853 | Tumor Luminal Proliferative | Endothelial |
| 7 | 2845 | Mesenchymal Progenitors | Tumor Basal EGFR |
| 8 | 2539 | Tumor Basal Active | Tumor Basal Active |
| 9 | 1966 | Fibroblast | Myoepithelial |
| 10 | 1378 | Tumor Luminal Proliferative | Macrophage |
| 11 | 679 | Macrophage | Plasma Cell |

**Table 4.**
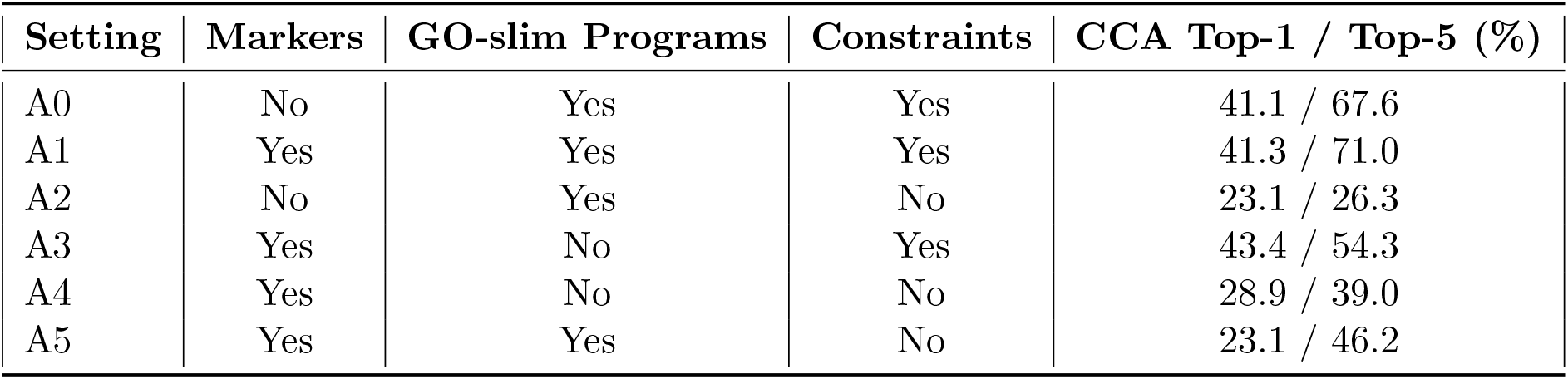
Ablation study evaluating NanoLLM label consistency using cell-weighted top-*k* evidence agreement. Results are reported for clusters with High or Moderate confidence (commitment = 86.57%). All settings share identical hSNMF clustering and marker identification; ablations vary only in evidence representation and label constraints.

In A0, cluster-level evidence is provided exclusively through GO-slim functional programs with supporting genes, without an explicit flat marker list. A1 corresponds to the full evidence configuration, in which both top-20 marker genes and GO-slim functional programs are provided under admissible label constraints. To isolate the effect of label constraints, we evaluate an unconstrained counterpart of A0 (A2), which removes label restrictions while retaining the same program-structured evidence. To assess the role of ontology-based functional structuring, we further consider marker-only configurations: A3 provides only top-20 marker genes under admissible label constraints, while A4 removes these constraints and allows open-ended label generation from marker genes alone. Finally, A5 removes admissible label constraints while retaining both marker genes and GO-slim functional programs, allowing the language model to generate labels without restriction despite having access to the full structured evidence representation.

#### Observations

Several trends emerge from the ablation analysis. First, admissible label constraints play a critical role in preventing unsupported predictions. Removing constraints substantially reduces agreement with evidence-supported labels. For example, Top-1 agreement drops from 41.3% in the full configuration (A1) to 23.1% when constraints are removed under program-only evidence (A2), and from 43.4% in the constrained marker-only setting (A3) to 28.9% in its unconstrained counterpart (A4). A similar degradation is observed in the full-evidence setting without constraints (A5), where Top-1 agreement drops from 41.3% in A1 to 23.1%, indicating that admissible label constraints remain important even when both markers and functional programs are provided.

Second, marker genes and ontology-derived functional programs provide complementary information. Under constrained inference, marker-only inputs (A3) yield the highest Top-1 agreement, reflecting stronger fine-grained specificity from marker genes. In contrast, program-structured inputs (A0) achieve higher Top-5 agreement, capturing lineage-consistent alternatives when subtype resolution is uncertain. The full evidence configuration (A1) combines these advantages, achieving the strongest overall Top-5 agreement.

Together, these results show that biological constraints and structured evidence jointly shape NanoLLM reasoning. Admissible label constraints reduce unsupported predictions, while marker-level and program-level evidence balance precision and robustness. This combination enables consistent cell-type annotation without reliance on labeled training data or supervised fine-tuning.

## 5 Conclusion

We presented NanoCellAnnotator, a biologically constrained and confidence-aware framework for automated cell-type annotation in spatial transcriptomics data. The method decouples spatial structure discovery, deterministic biological evidence construction, and language-model–based semantic inference, while restricting predictions to a curated admissible label space to prevent unsupported annotations.

A key feature of NanoCellAnnotator is explicit uncertainty handling through an independent confidence score derived from marker-gene support and lineage separation. This allows clusters with weak or conflicting molecular evidence to be flagged rather than forcibly assigned to predefined cell types.

Experiments on spatial transcriptomics datasets demonstrate that the framework can recover canonical cell populations while identifying clusters with ambiguous or mixed molecular signals. Ablation analyses further show that curated label constraints and structured biological evidence jointly contribute to robust annotation behavior.

Overall, NanoCellAnnotator provides a transparent and interpretable approach for confidence-aware cell-type annotation in spatial transcriptomics data.

## 6 Acknowledgments

This project was supported by the National Center for Advancing Translational Sciences (NCATS), National Institutes of Health, through Grant Award Number UM1TR004539. The content is solely the responsibility of the authors and does not necessarily represent the official views of the NIH.

## Supplementary Material

### Algorithm S1 Confidence-Based Ambiguity Handling (Cluster-Level)

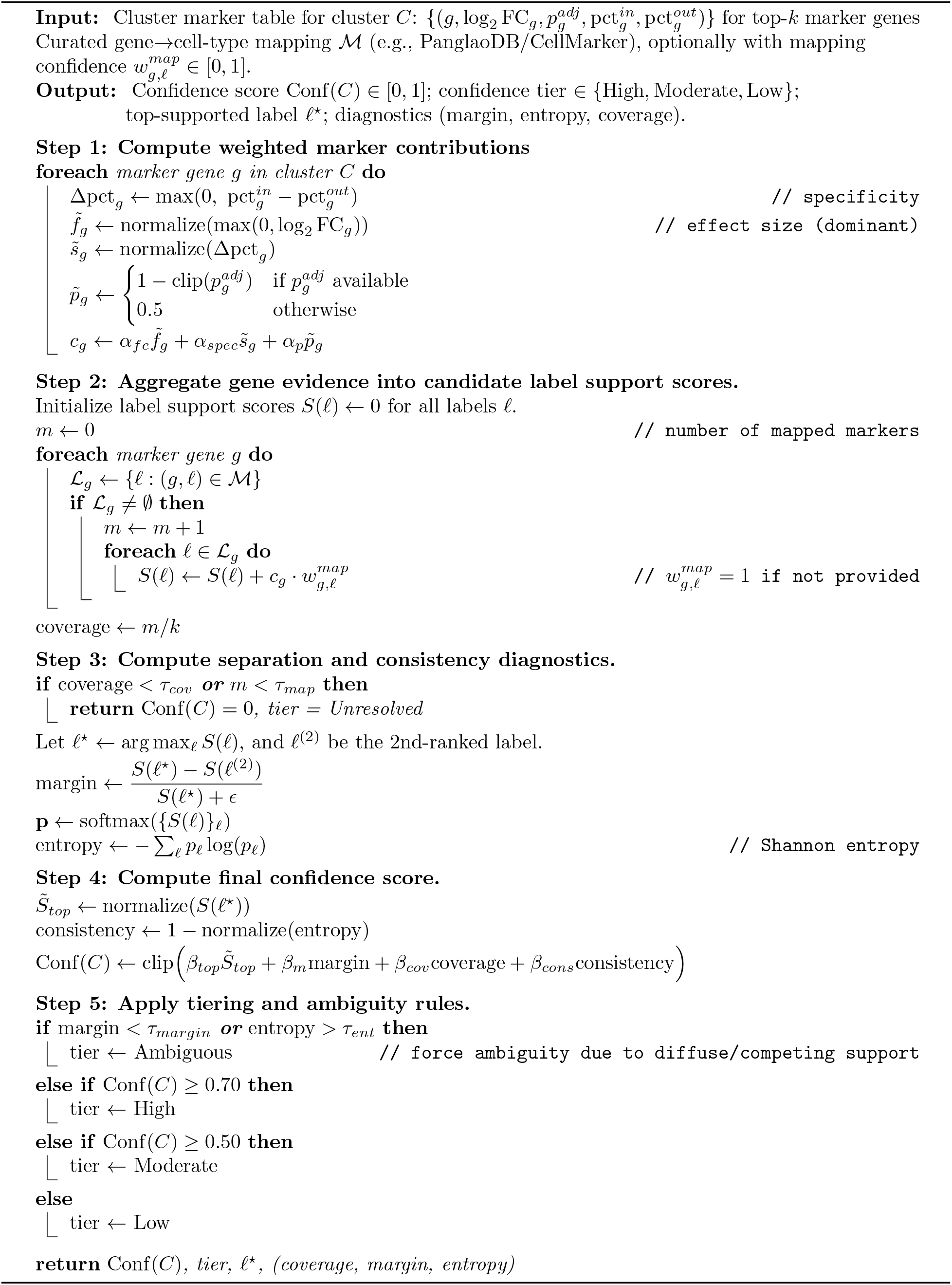

Algorithm S1 summarizes the deterministic procedure used to compute cluster-level confidence scores in NanoCellAnnotator. The algorithm aggregates marker-gene evidence into candidate cell-type support scores, evaluates separation between competing labels, and assigns confidence tiers based on evidence strength and consistency.

